## Supplemental Text 1 for "The impact of size on middle-ear sound transmission in elephants, the largest terrestrial mammal"

**S1 Text. Further details on elephant specimen preparation methods**

The elephant temporal bones arrived within variously sized portions of skull, depending on the age of the elephant and the equipment available for dissection at each zoo. Sizes of samples ranged from whole skulls for calves to a quarter of a skull for the adult (approximately 0.5 m^3^).

In all four elephant specimens, there was a thick cartilage-like tissue that divided the middle ear cavity into inferior and superior sub-cavities. The tissue spanned from lateral to medial bony walls of the middle ear cavity and attached to the malleus near the neck and had the tensor tympani muscle running through it. This tissue was first described in elephants by Buck [1] as the ‘tympanum proper,’ without any description of a potential function. It was also mentioned in Richards [2] as the membrane tegmen of the lower tympanum. We have not found any other reference to this tissue in the literature, except for a mention of potentially similar tissue in a turtle and in rare human pathology [3, 4]. Because the tympanum proper obstructed the malleus head and the entirety of the incus and stapes, we dissected this tissue out with a scalpel to access all three ossicles. Some initial umbo velocity measurements with this tissue intact showed that the removal of the tissue had little effect on the middle ear response, at least at the umbo. All results reported are with the tympanum proper removed.

When ETB6 and ETB7 were harvested from the elephant skull, an accidental cut above the ear canal removed the exterior bony wall but left the ear-canal soft tissue intact. Given that these specimens are difficult to obtain, we decided to repair them for use. A brass plate (ETB6) or piece of sturdy plastic (ETB7) shaped to fit the specimen was placed over the missing part of the bony ear canal and secured with dental cement. A 10 mm diameter brass ring was placed within the ear canal for ETB6 and ETB7 and secured with dental cement to allow for better placement of the sound source and probe-tube microphone positioning. They were positioned at the same depth in the ear canal as the samples that did not have the brass ring.

For ETB3 and ETB7, the promontory was damaged slightly during preparation and was repaired with dental cement. It is possible that air entered the cochlea, which would result in a lower cochlear input impedance (load to the middle ear) and likely higher stapes velocity at some frequencies. Since the results of these temporal bones were very similar to those in which the promontory was left intact (ETB2 and ETB6), we chose to include results from ETB3 and ETB7 since elephant specimens are rare.

Measurements were made during multiple sessions, each session lasting an entire day. This provided an opportunity to do test-retest measurements on each specimen. The temporal bone specimens were frozen and then thawed between repeated measurement sessions on different days. The thawing and refreezing of samples could potentially have contributed to degradation of tissue [5], or the draining of fluid from the cochlea [6]. However, measurements taken after several freeze-thaw sessions did not show significant changes in stapes velocity. The number of specimens for each ear type is too small to identify any possible age-, sex, or species-related trends.

To minimize effects due to drying, the temporal bones were continuously moistened using a room humidifier coupled to a tube placed near the opening of the MEC (Fig 1A). The humidifier generally remained on throughout a measurement session but was turned off during velocity measurements so as not to interfere with the laser beams of the 3D-LDV.

**References**

1.	Buck A.H. 1888 A Contribution to the Anatomy of the Elephant's Ear. (New Bedford, MA, Transactions of the American Otological Society.

2.	Richards H. 1890 *A further report on the anatomy of the elephant's ear.* New Bedford, MA, Mercury Publishing Company, Printers.

3.	Palva T., Northrop C., Ramsay H. 2001 Foreign body neonatal otitis media in infants. *Otol Neurotol* **22**(4), 433-443. (doi:<https://doi.org/10.1097/00129492-200107000-00003>).

4.	Palva T., Ramsay H., Böhling T. 1997 Tensor fold and anterior epitympanum. *Am J Otol* **18**(3), 307-316.

5.	Decraemer W.F., de La Rochefoucauld O., Funnell W.R., Olson E.S. 2014 Three-dimensional vibration of the malleus and incus in the living gerbil. *J Assoc Res Otolaryngol* **15**(4), 483-510. (doi:10.1007/s10162-014-0452-1).

6.	Ravicz M.E., Merchant S.N., Rosowski J.J. 2000 Effect of freezing and thawing on stapes-cochlear input impedance in human temporal bones. *Hear Res* **150**(1-2), 215-224. (doi:<https://doi.org/10.1016/s0378-5955(00)00200-8>).
