## Supplemental Text 2 for "The impact of size on middle-ear sound transmission in elephants, the largest terrestrial mammal"

**S2 Text. Measurements of 3D ossicle velocities**

A Polytec CLV-3D laser-Doppler vibrometer (3D LDV; Waldbronn, Germany) measured the velocity of discrete points. The three laser beams of the 3D LDV arranged in a cone pattern (cone half-angle 11.5 degrees) were reflected by a dichroic mirror, mounted to the 3D-LDV and motor assembly, and directed vertically downward and focused onto a reflective target on the elephant or human specimens (Fig 1A). The decoder unit of the 3D-LDV computed the resultant velocity of the target in the out-of-plane vertical (z) and in-plane (x and y) directions. The 3D-LDV was mounted on three-axis motorized and computer-controlled linear translation stages (6.25 µm resolution). This setup allowed the specimen to stay fixed while the 3D laser unit was moved to focus on the different reflective bead targets. Motion of the stages was controlled by SyncAV software (v0.34), which also generated the stimuli and recorded the synchronous responses. The coordinates of each measurement location were obtained and saved by SyncAv, allowing the 3D LDV to return to any of the previously measured locations.

The dichroic mirror mounted to the 3D-LDV allowed simultaneous visualization of the targets on the temporal bone through an operating microscope (Zeiss OPMI-1). A USB camera (DCC3240C; ThorLabs, Newton, NJ) connected to the microscope was used to monitor and capture images of the specimens. The camera was oriented vertically (along the Z axis) to take images of the X-Y plane. The dichroic mirror acts like a red-light filter. To improve the image quality through the microscope, the LDV assembly was moved away from the optical path and parked at a different location during ME manipulation, target placement, or image acquisition (e.g., Fig 4A).We specified a local “umbo” coordinate reference frame to describe malleus motion and a “stapes” coordinate reference frame to describe incus and stapes motions. These two anatomical coordinate reference frames differ and thus separate reference frames were required.

The sound source and probe-tube microphone assembly consisted of a loudspeaker (Vifa DPL28, Viborg, Denmark) coupled via a flexible tube to the inlet of a foam earplug (Etymotic, Elk Grove Village, IL) typically used as an insert earphone in clinical audiometry. The probe tube of an ER-7c (Etymotic) microphone was threaded through the foam plug. This earplug assembly was inserted into the ear canal opening and sealed as necessary with ear-canal-impression material (Silicast, Westone Laboratories, Colorodo Springs, CO).

Sound stimuli were generated and responses synchronously averaged using custom SyncAv software (version 0.42; [1]), which was developed using NI LabVIEW and PXI-4461 and PXI-4462 D/A and A/D hardware (NI, Texas). The stimuli were a series of pure tones generated in an 8192-sample buffer sampled at 30 kHz. For elephant, a series of 55 logarithmically spaced pure tones from 7 Hz to 13 kHz was used, and for human a series of 48 logarithmically spaced pure tones from 17 Hz to 11 kHz was used. For each tone, responses to 10 stimulus presentations were averaged together to improve the signal-to-noise ratio. The stimulus level was equalized as a function of frequency to provide an approximately constant ear-canal sound pressure (PEC) magnitude of approximately 90–100 dB SPL in elephant and 110 dB SPL in human. The noise floor was measured by repeating measurements with an output voltage close to 0 Volts. The PEC signal-to-noise ratio (SNR) was estimated to be typically more than 50 dB below 100 Hz, and at higher frequencies it was 60 dB or better.

Pieces of plastic tape (~0.2 mm^2^) with retroreflective glass-bead targets were affixed to the ossicular chain for velocity measurements: one on the umbo, three on other parts of the malleus, four on the incus, two on the stapes head, and one on each of the stapes crura (Fig 1A). Targets that shifted between measurement sessions were moved to approximately the same location as the prior session, based on digital images.

**References**

1.	Gottlieb P.K., Li X., Monfared A., Blevins N., Puria S. 2016 First results of a novel adjustable-length ossicular reconstruction prosthesis in temporal bones. *Laryngoscope* **126**(11), 2559-2564. (doi:10.1002/lary.25901).
