## Supplemental Text 3 for "The impact of size on middle-ear sound transmission in elephants, the largest terrestrial mammal"

**S3 Text. How piston directions were calculated**

The 3D LDV measures components of the velocity of the reflective target on an ossicle in three directions in the reference frame of the LDV: Vx and Vy in the horizontal plane/left-right and front-back, Vz in the vertical direction. Generally, the relevant directions of target velocity are not aligned with any of these axes. The pertinent velocities for describing ossicular motion functionally are aligned with anatomically relevant directions in an “intrinsic” reference frame (e.g., [1]). In the “umbo” coordinate system, the motion perpendicular to a plane approximately through the tympanic ring is termed “piston” motion of V_U_ and is assumed to be the most relevant motion for the tip of the malleus termed “umbo.” In the “stapes” coordinate system, the motion in a direction approximately perpendicular to the plane through the footplate and passing approximately through the stapes head is termed the stapes piston motion V_ST_ and is most relevant. The motion of the targets orthogonal to the piston direction (on the stapes head or posterior crus) are presumably due to rocking of the stapes about the footplate long or short axes and thought to not contribute appreciably to cochlear input [2, 3]. Since the elephant footplate is generally curved (Fig 1B), the motion of the stapes footplate plane become normal to the surface of the curved footplate, which effectively increases the volume velocity input the cochlea. The distal-tip of incus piston motion V_I_ was computed using the same local coordinate system as the stapes, assuming the same piston direction (Figs 1A and 4B).

Each bone was mounted in an orientation that enabled visibility of the reflective targets without interference by protruding bone and soft tissue of the specimen. Since each specimen varied in orientation, size and morphology (due to the age, species, and left or right ears), we transformed the velocities Vx, Vy, and Vz measured in the extrinsic coordinate system to anatomically appropriate ossicle-based velocities in an intrinsic coordinate system. To compute the motion in the relevant directions, the coordinate transformation between the LDV reference frame (extrinsic) and the relevant reference frame (intrinsic) must be computed. For example, for stapes velocities Vx, Vy, and Vz of the stapes target can be thought to have an elliptical motion trajectory with VMaj and VMin describing the ellipse. Vpist is the calculated projection of the ellipse along the piston direction.

The measured angles described above were used to project the measured Vx,y,z to the physiologically relevant “piston” umbo velocity V_U_, incus velocity V_I_ and stapes velocity V_ST_ using the Spatial Math Toolbox [4] in Matlab (Mathworks, Natick, MA). Because we measured the angles in a different order (Z-X-Y) than the Spatial Math Toolbox prefers, we recomputed the angles in the preferred Z-Y-X order – see S1 Table. The velocity of intermediate locations along the malleus (M2-M4) and incus (I2-I4) were computed in the same way described above.

In general, a coordinate transformation involves translation along the three axes and three rotations about axes. Translations have no effect on the velocity; it is only the rotations that must be computed. There are two conventions for describing rotations about axes: Euler, in which two rotations are applied about the same axis, and Cardinian, in which there is a rotation about each axis. We chose to use Cardinian rotations, which are commonly used in aeronautics and robotics (e.g., [4]) and which we found more intuitive. For example, to transform the measured stapes x, y, and z velocities to the intrinsic stapes reference frame, the extrinsic LDV reference frame may need to be rotated about one, two, or all three of the x-y-z axes (see S1 Fig).

The transformation from the measured reference frame to the intrinsic reference frame can be described easily in a matrix representation. For example, a velocity Vx,y,z measured in the extrinsic (LDV) reference frame has components Vx, Vy, and Vz in the x-, y-, and z-directions, respectively, and can be represented as a vector:

$V_{xyz}= \left\{ \begin{matrix} V_{x} \\ V_{y} \\ V_{z} \end{matrix} \right\}.$	(Eq. S 1)

The transformation of this velocity into a new reference frame V_abc_ rotated through an angle $\theta$ about the z axis can be described by a rotation matrix Rz to multiply Vxyz by:

$V_{abc} about z= \left\{ \begin{matrix} V_{a} \\ V_{b} \\ V_{c} \end{matrix} \right\}= \left[ \begin{matrix} \cos\theta& -sin \theta& 0 \\ \sin\theta& \cos\theta& 0 \\ 0 & 0 & 1 \end{matrix} \right]\left\{ \begin{matrix} V_{x} \\ V_{y} \\ V_{z} \end{matrix} \right\}= R_{z}\left\{ \begin{matrix} V_{x} \\ V_{y} \\ V_{z} \end{matrix} \right\}.$	(Eq. S 2)

Similarly, rotations about the x or y axes can be described by rotation matrices Rx or Ry:

$V_{abc} about x= \left[ \begin{matrix} 1 & 0 & 0 \\ 0 & \cos\theta& -sin \theta\\ 0 & \sin\theta& \cos\theta\end{matrix} \right]\left\{ \begin{matrix} V_{x} \\ V_{y} \\ V_{z} \end{matrix} \right\}= R_{x}\left\{ \begin{matrix} V_{x} \\ V_{y} \\ V_{z} \end{matrix} \right\}$ , and	*(Eq. S 3)*

$V_{abc} about y= \left[ \begin{matrix} \cos\theta& 0 & \sin\theta\\ 0 & 1 & 0 \\ -sin \theta& 0 & \cos\theta\end{matrix} \right]\left\{ \begin{matrix} V_{x} \\ V_{y} \\ V_{z} \end{matrix} \right\}= R_{y}\left\{ \begin{matrix} V_{x} \\ V_{y} \\ V_{z} \end{matrix} \right\}$ .	*(Eq. S 4)*

Then successive rotations about the x, (new) y, and (new) z axes can be computed by multiplying the rotation matrices together (from right to left). For example, to obtain velocities V_Maj_, V_pist_, V_min_ in the Major axis – piston – minor axes of the stapes intrinsic reference frame:

$V_{Maj-pist-min}= \left\{ \begin{matrix} V_{Maj} \\ V_{pist} \\ V_{min} \end{matrix} \right\}= R_{z}\times R_{y}{\times R}_{x}\left\{ \begin{matrix} V_{x} \\ V_{y} \\ V_{z} \end{matrix} \right\}= R_{zyx}\left\{ \begin{matrix} V_{x} \\ V_{y} \\ V_{z} \end{matrix} \right\}$(Eq. S 5)

(where $\times$ signifies matrix multiplication; e.g., [4]).

When applying rotations, the order in which the rotations are applied matters (rotational displacements are not commutative – e.g., [4]). In addition, rotations about some axes are easier to measure than others. For these experiments, we found that azimuth (rotation about z, of the projection of the piston axis onto the x-y plane) was the easiest to measure, then elevation (rotation about x, angle of the piston axis from the x-y plane) – see S3 Fig. Rotation about y (to orient the stapes footplate major axis along the x axis) was most difficult to measure, was generally an iterative process, and had the least influence on the computation of piston-direction motion. See S1 Table for measured azimuth and elevation and estimated y rotation angles for both stapes piston direction and for the “umbo” direction (perpendicular to the tympanic ring).

S3 Fig shows plots of the rotations measured for human temporal bone TB20, superposed over a photograph of the stapes and surroundings taken in the x-y plane/-z direction taken by the camera through the dichroic mirror on the 3D LDV. The rotations were applied sequentially in z-x-y order as they were measured. We rotated the 3D plot in MATLAB to check that the projected axes matched the anatomy. Panel A shows the x-y-z (extrinsic) coordinate system centered approximately at the center of the stapes footplate (out of view); panel B shows a rotation of the *x* and *y* axes (shown by thicker lines) +20 degrees (azimuth) about the *z* axis; panel C shows an additional rotation of the (new) *y* and *z* axes (thicker lines) of +40 degrees about the (new) *x* axis (intrinsic Maj axis), and panel E shows an additional rotation of the new *x* and *z* axes of –150 degrees about the (new) *y* axis (intrinsic pist axis) to arrive at the new Maj-pist-min intrinsic reference frame.

To perform the coordinate transformations, we used the Spatial Math Toolbox [4] in MATLAB. The Spatial Math Toolbox performs the coordinate transformation rotations in the order z-y-x, while we measured them in the order z-x-y. We used the Spatial Math Toolbox function torpy to compute the corresponding x, y, and z rotations to be applied in the order z-y-x (S1 Table). In the case that one or more of azimuth, elevation, or y rotation is 0 (as for all umbo direction transformations), the rotation angles are unchanged but may appear in a different order. When all of azimuth, elevation, and y rotation are non-zero (as for TB20 SFP), the x, y, and z rotations may be quite different from the measured angles (S3 Fig), but the corresponding transformation matrix is the same.

We checked the transformation matrix by computing the new projected Major, piston, and minor axes and superposing them on photos of the temporal bones in the x-y plane (S3 Fig). Panel E shows the extrinsic coordinate system as before; panel F applies a –139-degree rotation about the *z* axis; panel G then applies a –22-degree rotation about the (new) *y* axis; and panel H finally applies a +135-degree rotation about the (new) *x* axis. It can be seen easily that applying different rotations in a different order produces the same result: the intrinsic coordinate system.

**References**

1.	Decraemer W.F., de La Rochefoucauld O., Dong W., Khanna S.M., Dirckx J.J., Olson E.S. 2007 Scala vestibuli pressure and three-dimensional stapes velocity measured in direct succession in gerbil. *J Acoust Soc Am* **121**(5 Pt1), 2774-2791. (doi:<https://doi.org/10.1121/1.2709843>).

2.	Huber A.M., Sequeira D., Breuninger C., Eiber A. 2008 The effects of complex stapes motion on the response of the cochlea. *Otol Neurotol* **29**(8), 1187-1192. (doi:<https://doi.org/10.1097/MAO.0b013e31817ef49b>).

3.	Eiber A., Huber A.M., Lauxmann M., Chatzimichalis M., Sequeira D., Sim J.H. 2012 Contribution of complex stapes motion to cochlea activation. *Hear Res* **284**(1-2), 82-92. (doi:<https://doi.org/10.1016/j.heares.2011.11.008>).

4.	Corke P.I. 2017 Robotics, Vision \& Control: Fundamental Algorithms in MATLAB. (Springer).
