## Supplemental Text 4 for "The impact of size on middle-ear sound transmission in elephants, the largest terrestrial mammal"

**S4 Text. Computation of velocities in anatomically relevant piston directions**

A 1D LDV has traditionally been used for characterizing sound driven middle ear (ME) motion responses. For measurements in human temporal bones, the stapes footplate was accessed with an LDV beam angled relative to the desired stapes piston direction. This angle varied depending on the facial recess opening but was in the 30 to 45 degrees range. The cosine of the angle was typically used to approximate the motion in the piston direction. Though such corrections may only be accurate at lower frequencies [1], the present transformation methods should prove to be accurate at all frequencies.

For umbo measurements, the LDV beam angle was typically from the ear canal approach, which was nearly perpendicular to the plane of the ear drum (tympanic membrane or TM). Umbo measurements have also been made through the posterior-inferior ME opening. These measurements were from an oblique angle rather than from the desired direction orthogonal to the plane of the TM.

Fig 2B and 2D shows V_ST_ measurements in human temporal bones using both 1D and 3D LDV methods are generally in good agreement, but with some important differences. For the Aibara et al. [2] measurements, the lowest frequency reported was 50 Hz whereas the present measurements go down to about 17 Hz. Raufer et al. [3] reported the velocity of the stapes posterior crus from 2 kHz down to the infrasound region with the lowest frequency of 0.9 Hz. These measurements of V_ST_ were made with a 1D LDV from approximately a 30-degree angle from the piston direction, and a cosine correction was applied.

Below about 200 Hz, our measurements are lower in magnitude and steeper than those of Aibara et al. [2] and Raufer et al. [3] and the phase is generally a bit higher (but within about +¼ cycle). For both the present and Raufer et al. [3] studies, measurements were made with a vibration isolation table while such a table was not available to Aibara et al. [2]. This might explain the higher measured magnitudes in Aibara et al. [2] below 120 Hz.

Above the human ME resonant frequency of about 1 kHz, the high frequency slope for the Aibara et al. [2] stapes magnitude is a bit steeper (about -6 dB/oct) than the present measurements (about -5 dB/oct) while the phases nearly overlap. Also, above 500 Hz, the Raufer et al. [3] magnitude is lower by 10 dB or more and the phase decreases more rapidly than Aibara et al. [2] and present results. It may be that these differences occur due to measurement location difference. Aibara et al. [2] made measurements of the stapes footplate, which is accessible by a 1D LDV. However, the footplate was not accessible to the three beams of our 3D-LDV, so we measured the velocity of stapes posterior crus when accessible.

Similar 3D motion studies have been performed in the gerbil [4] and human [5]. A Guinea pig study showed that energy going into the cochlea was a combination of both the rotational motion and piston direction motion of the stapes that contributed to energy entering the cochlea [6]. Our 3D LDV results suggest that the elephant might also have multiple modes of motion involved in the transfer function.

The transformation of 3D-LDV **V_X,Y,Z_** measurements of each target from the measured extrinsic LDV coordinate system to the intrinsic ossicle coordinate system was done by first estimating a series of angles required to rotate the extrinsic coordinate system to the intrinsic coordinate system. We measured: (a) the angle between the projection of the malleus or stapes “piston” direction onto the 3D LDV X-Y plane (looking vertically down through the camera) and the Y axis (azimuth = rotation about 3D LDV Z axis); (b) the angle between the malleus or stapes “piston” direction and the X-Y plane (elevation = rotation about (rotated) 3D-LDV X axis); and (c) the angle of rotation necessary about the piston axis to orient the stapes footplate major axis in the posterior direction (roll = rotation about (rotated) 3D LDV Y axis). These angles are illustrated in S3 Fig. For ETB6 and ETB7 angle measurements were performed in situ and checked against photos taken by the 3D LDV camera, while for ETB2, and ETB3 and human specimens, angles were estimated from photos only (S1 Table lists the series of angles measured). The umbo coordinate system was determined through appropriate angle transformations. The measured LDV coordinate reference frame to anatomical piston direction coordinate frame is illustrated for the stapes in S1A Fig and for the umbo in S1B Fig.

**References**

1.	Chien W., Ravicz M.E., Merchant S.N., Rosowski J.J. 2006 The effect of methodological differences in the measurement of stapes motion in live and cadaver ears. *Audiol Neurootol* **11**(3), 183-197. (doi:<https://doi.org/10.1159/000091815>).

2.	Aibara R., Welsh J.T., Puria S., Goode R.L. 2001 Human middle-ear sound transfer function and cochlear input impedance. *Hearing Research* **152**, 100-109. (doi:10.1016/s0378-5955(00)00240-9).

3.	Raufer S., Masud S.F., Nakajima H.H. 2018 Infrasound transmission in the human ear: Implications for acoustic and vestibular responses of the normal and dehiscent inner ear. *J Acoust Soc Am* **144**(1), 332. (doi:<https://doi.org/10.1121/1.5046523>).

4.	Decraemer W.F., de La Rochefoucauld O., Funnell W.R., Olson E.S. 2014 Three-dimensional vibration of the malleus and incus in the living gerbil. *J Assoc Res Otolaryngol* **15**(4), 483-510. (doi:10.1007/s10162-014-0452-1).

5.	Dobrev I., Ihrle S., Roosli C., Gerig R., Eiber A., Huber A.M., Sim J.H. 2016 A method to measure sound transmission via the malleus-incus complex. *Hear Res* **340**, 89-98. (doi:<https://doi.org/10.1016/j.heares.2015.10.016>).

6.	Eiber A., Huber A.M., Lauxmann M., Chatzimichalis M., Sequeira D., Sim J.H. 2012 Contribution of complex stapes motion to cochlea activation. *Hear Res* **284**(1-2), 82-92. (doi:<https://doi.org/10.1016/j.heares.2011.11.008>).
