## Supplemental Code 1 for "The impact of size on middle-ear sound transmission in elephants, the largest terrestrial mammal": S1_Code_OCRetal_Ele_AC_bioRxiv_submittedPLUSONE_09252023.Rmd

````
---
Title: "S1 Code for 'The impact of size on middle-ear sound transmission in elephants, the largest
  terrestrial mammal'"
Authors: "Caitlin E. O'Connell-Rodwell, Jodie Berezin, Anbuselvan Dharmarajan, Michael
  E. Ravicz, Yihan Hu, Xiying Guan, Kevin O’Connor, Sunil Puria"
Most recently updated: August 25th, 2023
output: pdf_document
---

The script below can be used to replicate the statistical analysis using the S2 Data.

Questions regarding the data or scripts can be directed to the corresponding author, Caitlin E. O'Connell-Rodwell at caitlin_oconnell[at]meei.harvard.edu


*Table of Contents*
1. Data and Model Descriptions (pre-code information)
2. Loading Libraries and Importing Data
3. Data Wrangling
4. Modeling
5. Supplemental: Raw data modeling (for model assumption violation details)


*1. Data and Model Descriptions*

**Eight mixed-effects models were used for the statistical analysis:**
1) Comparing the elephant and human magnitude for umbo and stapes velocity combined
2) Comparing the elephant and human phase for umbo and stapes combined
3) Comparing the elephant and human magnitude for umbo velocity
4) Comapring the elephant and human phase for umbo velocity
5) Comparing the elephant and human magnitude for stapes velocity
6) Comparing the elephant and human phase for stapes velocity
7) Comparing elephant umbo and stapes magnitude
8) Comparing human umbo and stapes magnitude

**Data Description**
Mean velocity data for each specimen was extracted from Figure S2. Details below are the measurement ID number and n measurements collected for each specimen which make up the mean (see folder containing these data files). The number in parantheses next to each velocity measurement represents the number of tones logarithmically spaced in the sound stimuli for each mean (per specimen), from 17 Hz to 11000 Hz (see Methods section for more information).

    *Umbo*                      *Stapes*
ETB2: 7008 (50)             ETB2: 7002,7004 (50)
ETB3: 3002,5002 (50)        ETB3: 5012,5013 (50)
ETB6: 1012,2002 (50)        ETB6: NA
ETB7: 1002,2022,2030 (50)   ETB7: 1024,1027,2003,2015,2032 (50)
TB18: NA                    TB18: 1007,1008 (48)
TB19: 1012 (48)             TB19: 1004,1005 (48)
TB20: 1067 (48)             TB20: 1058,1059 (48)


For all models, frequency is in Hz.

Alpha set to 0.05 for all models.


# 2. Loading Libraries, Importing Data

## Set the working directory
```{r}
#setwd()
```

## load the packages used
```{r}
library(rstatix) # checking model fits
library(lmerTest) # running models
library(tidyverse) # data wrangling
```

## Load the data
```{r}
library(readr)
velocity <- read_csv("O'Connell-Rodwelletal_RoySoc_VelocityData_StatsAnalysis_Aug2023.csv")
View(velocity)

str(velocity)
```

# 3. Data Wrangling

## Change everything to factors and numeric variables. 

The "as.character" function within the factor functions keeps the name of the factors when models are ran.
```{r}
velocity$Group <- as.factor(as.character(velocity$Group))
velocity$Specimen <- as.factor(as.character(velocity$Specimen))
velocity$Ossicle <- as.factor(as.character(velocity$Ossicle))

velocity$Freq_x <- as.numeric(velocity$Freq_x)
velocity$Mag_y <- as.numeric(velocity$Mag_y)
velocity$Phase_y <- as.numeric(velocity$Phase_y)
```


## Make a new category with low and high frequencies

Note: For direct comparisons, all frequencies below 17 Hz (the minimum human frequency collected) and above 11 kHz (the maximum frequency collected for humans) were removed. Four categories were created: "below17" (frequencies below 17 Hz), "low" (frequencies from 17 Hz to 1000 Hz), "high" (frequencies > 1000 Hz and <= 11 kHz), and "above11kHz" (frequencies > 11 kHz).

```{r}
velocity2 <- velocity %>%
  mutate(freq_cat = ifelse(Freq_x < 17, "below17",
           ifelse(Freq_x >= 17 & Freq_x <= 1000, "low", 
                  ifelse(Freq_x > 1000 & Freq_x <= 11000, "high",
                         ifelse(Freq_x > 11000, "above11kHz", NA)))))

velocity2$freq_cat <- as.factor(as.character(velocity2$freq_cat))
levels(velocity2$freq_cat) # check it

velocity3 <- velocity2 %>%
  filter(freq_cat != "below17") %>% # removing frequencies lower than 14 Hz
  filter(freq_cat != "above11kHz") %>% # removing frequencies higher than 11 kHz
  droplevels() # removes the dropped factor categories when checking the categories for velocity3$freq_cat

levels(velocity3$freq_cat) # check it again
  
```

## Transforming the data

- All models using raw data violated normality and homogeneity of variance of assumptions (see section 5 for details).
- Magnitude can be log transformed.
- Due to negative values (lowest value -3.49), the phase data was transformed by taking the log of the phase plus four: log(phase + 4).

```{r}
velocity4 <- velocity3 %>%
  mutate(Log_Mag_y = log(Mag_y)) %>%
  mutate(Log_Phase_y = log(Phase_y + 4))
```

## Filtering on ossicle for models 3, 4, 5, and 6

### Filter for umbo data
```{r}
velocity_umbo <- velocity4 %>%
  filter(Ossicle == "Umbo")
```

### Filter for stapes data
```{r}
velocity_stapes <- velocity4 %>%
  filter(Ossicle == "Stapes")
```


# 4. Modeling

## Model 1: Comparing the elephant and human magnitude for umbo and stapes velocity combined.

### Running the model
```{r}
model1 <- lmer(Log_Mag_y ~ Group*Ossicle + (1|Freq_x/Specimen), data = velocity4)

summary(model1)
```

### Checking the model fit
```{r}
qqnorm(resid(model1))
qqline(resid(model1))

plot(resid(model1))
abline(h = 0, col = "blue")
```


## Model 2: Comparing the elephant and human phase for umbo and stapes combined.

### Running the model
```{r}
model2 <- lmer(Log_Phase_y ~ Group*Ossicle + (1|Freq_x/Specimen), data = velocity4)

summary(model2)
```

### Checking the model fit
```{r}
qqnorm(resid(model2))
qqline(resid(model2))

plot(residuals(model2))
abline(h = 0, col = "blue")
```


## Model 3: Comparing the elephant and human magnitude for umbo velocity.

### Running the model
```{r}
model3 <- lmer(Log_Mag_y ~ Group*freq_cat + (1|Specimen), data = velocity_umbo)

summary(model3)
```

### Checking the model fit
```{r}
qqnorm(resid(model3))
qqline(resid(model3))

plot(residuals(model3))
abline(h = 0, col = "blue")
```


## Model 4: Comapring the elephant and human phase for umbo velocity. 

### Running the model
```{r}
model4 <- lmer(Log_Phase_y ~ Group*freq_cat + (1|Specimen), data = velocity_umbo)

summary(model4)
```

### Checking the model fit
```{r}
qqnorm(resid(model4))
qqline(resid(model4))

plot(residuals(model4))
abline(h = 0, col = "blue")
```


## Model 5: Comparing the elephant and human magnitude for stapes velocity.

### Running the model
```{r}
model5 <- lmer(Log_Mag_y ~ Group*freq_cat + (1|Specimen), data = velocity_stapes)

summary(model5)
```

### Checking the model fit
```{r}
qqnorm(resid(model5))
qqline(resid(model5))

plot(residuals(model5))
abline(h = 0, col = "blue")
```


## Model 6: Comparing the elephant and human phase for stapes velocity.

### Running the model
```{r}
model6 <- lmer(Log_Phase_y ~ Group*freq_cat + (1|Specimen), data = velocity_stapes)

summary(model6)
```

### Checking the model fit
```{r}
qqnorm(resid(model6))
qqline(resid(model6))

plot(residuals(model6))
abline(h = 0, col = "blue")
```


## Model 7: Comparing elephant umbo and stapes magnitude

### filtering the data
```{r}
elephant.only <- velocity4 %>%
  filter(Group == "Elephant")
```

### Running the model
```{r}
model7 <- lmer(Log_Mag_y ~ Ossicle*freq_cat + (1|Specimen), data = elephant.only)

summary(model7)
```

### Checking the model fit
```{r}
qqnorm(resid(model7))
qqline(resid(model7))

plot(residuals(model7))
abline(h = 0, col = "blue")
```


## Model 8: Comparing human umbo and stapes magnitude

### Filter the data
```{r}
human.only <- velocity4 %>%
  filter(Group == "Human")
```

### Running the model
```{r}
model8 <- lmer(Log_Mag_y ~ Ossicle*freq_cat + (1|Specimen), data = human.only)

summary(model8)
```

### Checking the model fit
```{r}
# residuals
qqnorm(resid(model8))
qqline(resid(model8))

plot(residuals(model8))
abline(h = 0, col = "blue")
```


# 5. Supplemental: Raw data modeling (for model assumption violation details)

Note: These model results were not included in the manuscript. These models are just to display the model assumption violations when raw data was used.


## Model 1.2: Comparing the elephant and human magnitude for umbo and stapes velocity combined.

### Running the model
```{r}
model1.2 <- lmer(Mag_y ~ Group + Ossicle + (1|Freq_x/Specimen), data = velocity4)

summary(model1.2)
```

### Checking the model fit
```{r}
qqnorm(resid(model1.2))
qqline(resid(model1.2))

plot(resid(model1.2))
abline(h = 0, col = "blue")
```


## Model 2.2: Comparing the elephant and human phase for umbo and stapes combined.

### Running the model
```{r}
model2.2 <- lmer(Phase_y ~ Group*Ossicle + (1|Freq_x/Specimen), data = velocity4)

summary(model2.2)
```

### Checking the model fit
```{r}
qqnorm(resid(model2.2))
qqline(resid(model2.2))

plot(residuals(model2.2))
abline(h = 0, col = "blue")
```


## Model 3.2: Comparing the elephant and human magnitude for umbo velocity.

### Running the model
```{r}
model3.2 <- lmer(Mag_y ~ Group*freq_cat + (1|Specimen), data = velocity_umbo)

summary(model3.2)
```

### Checking the model fit
```{r}
qqnorm(resid(model3.2))
qqline(resid(model3.2))

plot(residuals(model3.2))
abline(h = 0, col = "blue")
```


## Model 4.2: Comparing the elephant and human phase for umbo velocity.

### Running the model
```{r}
model4.2 <- lmer(Phase_y ~ Group*freq_cat + (1|Specimen), data = velocity_umbo)

summary(model4.2)
```

### Checking the model fit
```{r}
qqnorm(resid(model4.2))
qqline(resid(model4.2))

plot(residuals(model4.2))
abline(h = 0, col = "blue")
```


## Model 5.2: Comparing the elephant and human magnitude for stapes velocity.

### Running the model
```{r}
model5.2 <- lmer(Mag_y ~ Group*freq_cat + (1|Specimen), data = velocity_stapes)

summary(model5.2)
```

### Checking the model fit
```{r}
qqnorm(resid(model5.2))
qqline(resid(model5.2))

plot(residuals(model5.2))
abline(h = 0, col = "blue")
```


## Model 6.2: Comparing the elephant and human phase for stapes velocity.

### Running the model
```{r}
model6.2 <- lmer(Phase_y ~ Group*freq_cat + (1|Specimen), data = velocity_stapes)

summary(model6.2)
```

### Checking the model fit
```{r}
qqnorm(resid(model6.2))
qqline(resid(model6.2))

plot(residuals(model6.2))
abline(h = 0, col = "blue")
```


## Model 7.2: Comparing elephant umbo and stapes magnitude

### Running the model
```{r}
model7.2 <- lmer(Mag_y ~ Ossicle*freq_cat + (1|Specimen), data = elephant.only)

summary(model7.2)
```

### Checking the model fit
```{r}
qqnorm(resid(model7.2))
qqline(resid(model7.2))

plot(residuals(model7.2))
abline(h = 0, col = "blue")
```


## Model 8.2: Comparing human umbo and stapes magnitude

### Running the model
```{r}
model8.2 <- lmer(Mag_y ~ Ossicle*freq_cat + (1|Specimen), data = human.only)

summary(model8.2)
```

### Checking the model fit
```{r}
qqnorm(resid(model8.2))
qqline(resid(model8.2))

plot(residuals(model8.2))
abline(h = 0, col = "blue")
```
````
